# Integration of single-cell multi-omic data with graph-based topic modelling

**DOI:** 10.64898/2026.02.25.707947

**Authors:** Gabriele Malagoli, Filippo Valle, Andreina Tirabassi, Annalisa Marsico, Loredana Martignetti, Michele Caselle, Maria Colomé-Tatché

## Abstract

Recent advances in single-cell biology enable the profiling of multiple molecular layers, such as the transcriptome, epigenome, and surface proteins, within a single cell. Tackling the complexity of these data from different perspectives allows researchers to get the most complete insights into the biological properties of cells. Here, we propose a graph-based topic modelling method called bionSBM. Our method leverages well-known community-detection methods for multipartite graphs and the interpretability of topic modelling to cluster and explain high-dimensional, sparse, and noisy single-cell matrices. We applied our algorithm to paired single-cell multi-omics data, such as 10X Multiome, SHARE-seq, and CITE-seq. We showed that it achieves superior performance compared to state-of-the-art methods for ground-truth label retrieval, with high specificity and distinct biological interpretability.

## Background

The emergent phenotypes of biological systems result from complex molecular interactions. These interactions play a role at multiple levels of gene expression regulation: genome, epigenome, transcriptome and proteome. Characterising such regulatory behaviour is crucial to studying these processes. It is an essential step towards the development of drugs and targeted personalised medicine strategies[1], as well as for other translational applications such as drug discovery and validation[2] or tumour microenvironment profiling. The latest advances in single-cell sequencing technologies, such as 10X Multiome and SHARE-seq[3], enable profiling of gene expression and chromatin accessibility at the single-cell level. Moreover, technologies such as CITE-seq[4] allow profiling not only gene expression but also the measurement of surface molecules using antibody-derived tags (ADTs). All these techniques are opening new avenues towards the comprehension of complex gene regulatory mechanisms[5].

The development of experimental protocols poses a new challenge for the scientific community: algorithms and downstream pipelines must be adapted to integrate paired and multi-feature data. In particular, when analyzing these data, one is typically interested in their modular structure. The main goal is to identify groups of genes and genomic loci that act in a coordinated way, and groups of cells with similar molecular patterns. Given the multimodal nature of the data, the focus can be on groups of genes, ATAC-seq peaks, ADTs, or any features the measurements represent. Moreover, it is of interest to link these modular structures across the omic layers and use these modules to infer clusters within cells, which is a highly non-trivial task. Several algorithms have been proposed in the past few years to address it. A popular approach is deep learning. Deep learning methods, such as autoencoders[6], perform effective dimensionality reduction, but they are sensitive to differences in input scale, which is a key characteristic of multimodal data[7]. To overcome these limitations, here we will concentrate on a class of algorithms based on topic modelling. Topic modelling algorithms were initially developed in Natural Language Processing (NLP) for the automatic identification of “topics” in a corpus of texts, using only the frequency of words as input. They can group sets of words that “speak about the same theme” and, in this way, identify the texts’ topics. There is an obvious analogy between this task and the one we are interested in. If we map texts to cells and the frequency of words to gene expression levels, ATAC-seq peak openness/ADT levels, the same approach described above can be used to identify and characterise cell populations based on their patterns of molecular measurements. In this analogy, topics are groups of genes/open genomics loci/surface proteins that act cooperatively, and these regulomes and pathways can be used to characterize different cell populations. The rationale for using topic modelling methods for single-cell data is that the transcriptome of cells exhibits statistical properties similar to those of words in texts[8].

One of the most interesting features of topic modelling algorithms is that they output probability distributions over cells belonging to each topic, which better describe the complexity of biological systems than a naive, deterministic association in which cells are simply assigned to a group (or cluster). The topics, on the other hand, are not only lists of loci; they are described as probability distributions over all genomic features.

The most popular topic models are either based on Latent Dirichlet Allocation (LDA)[9], Non-Negative Matrix factorisation (NMF)[10] or Stochastic Block Models (SBM)[11]. LDA approaches the problem of topic modelling by assuming a Dirichlet prior for the distribution of words within topics and of topics within documents (in our setting, corresponding to the distributions of genes, peaks, and ADTs within topics, and of topics within cells). The choice of this prior does not have a biological foundation; it simply provides a convenient implementation to solve the topic modelling problem[12]. Since the Dirichlet distribution is unimodal, regardless of the choice of hyperparameters, it assumes a single shape for the distribution of words into topics and topics into documents, which is a strong homogeneity assumption difficult to reconcile with the heterogeneity of biological systems[13]. Another common approach is Nonnegative Matrix Factorisation (NMF)[14], which is also very effective computationally. However, it has two main limitations: its linearity limits the ability to understand complex interactions, which are typical in biological systems, and it tends to find steep boundaries among clusters, which can be a problem when dealing with highly heterogeneous data, as is the case in single-cell sequencing[12]. We will therefore concentrate on the following in the third category of topic models, based on Stochastic Block Modelling (SBM)[11,13,15]. We present a graph-based method, bionSBM, for integrating and interpreting paired single-cell multi-omic data [Fig. 1]. bionSBM can cluster cells based on any modality a user needs to consider (peaks, ADTs, mRNAs, the complete transcriptome, etc.), treating them equally. It does not impose a unimodal prior but defines an agnostic prior to better fit the topics-cells and features-topics distributions. This algorithm is part of a long-standing effort to apply SBM methods to biomedical data, which started with the construction of a hierarchical version of SBM based topic modeling (hSBM)[12] for gene expression data, then expanded to bulk tissue multiomics data (nSBM)[16], to single cell gene expression data[17] and finally in the present paper to single cell multiomics data. In the following, we first describe the main features of bionSBM and then compare it with two other topic modelling algorithms for single-cell data analysis, ShareTopic[18] and Mowgli[19], which are state-of-the-art algorithms for multimodal single-cell analysis based, respectively, on LDA and NMF.

**Figure 1.**
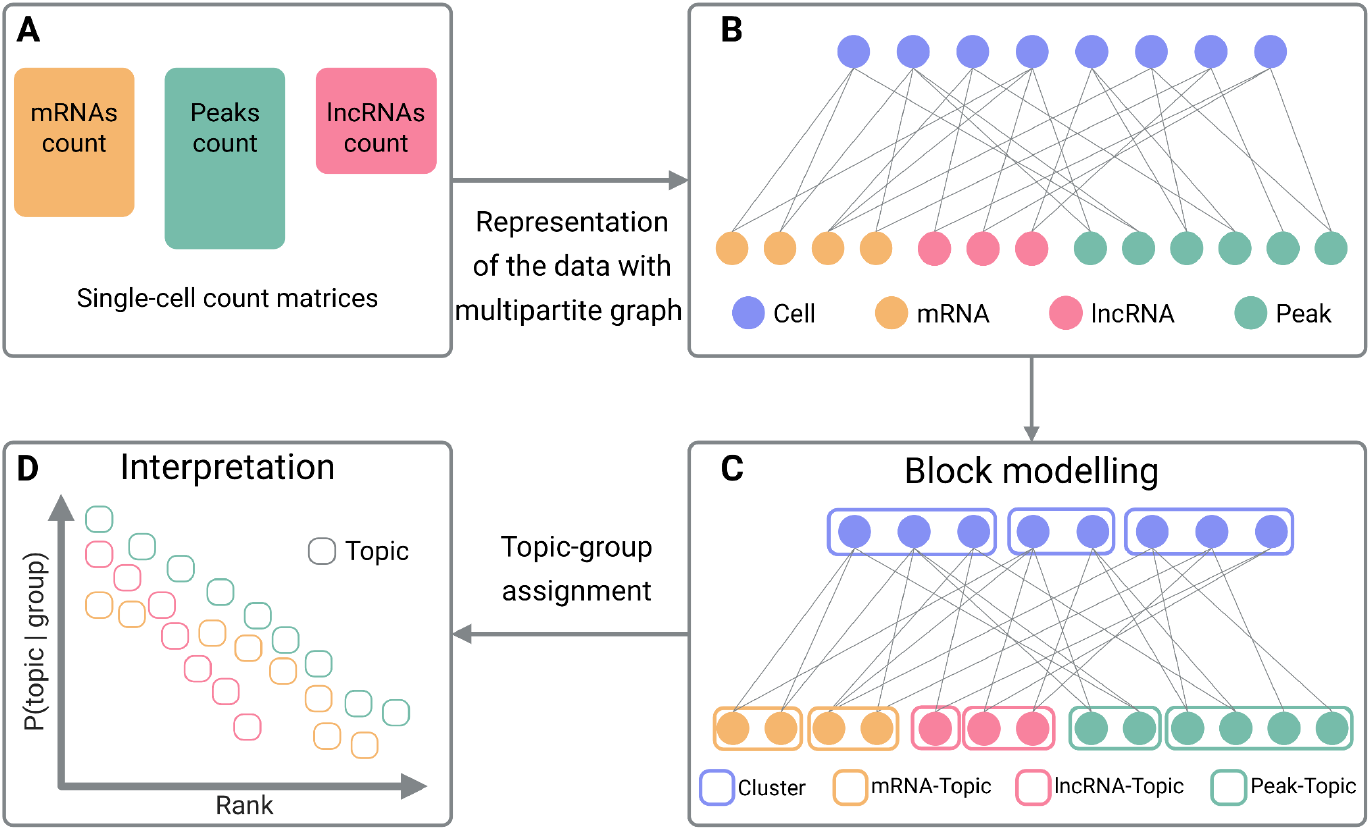
schematic pipeline of the bionSBM algorithm. (**A**) Data is input as count matrices. (**B**) Input matrices are used to build a multipartite network: features are connected to the cell branch if the feature is expressed / open in the cell with weighted edges, but features are not connected. (**C**) bionSBM finds the best block model to group the cells into clusters and the features into topics. (**D**) The explanation for the cell grouping can then be easily obtained by using the topic distribution probabilities within clusters.

## Results

### The bionSBM model

bionSBM leverages a class of algorithms called Stochastic Block Models (SBMs)[11]. In particular, bionSBM is based on a particularly efficient hierarchical version of SBM developed in[13,15], known as hierarchical Stochastic Block Modelling (hSBM), which was later extended to n-partite SBM (nSBM) for multipartite graphs.

The idea behind hSBM is to maximize the posterior probability of the model given the data, namely. *P* (*model* | *data*). In a Bayesian framework, this is done by assuming a model *P* (*model*) and maximizing the likelihood *P* (*model* | *data*). hSBM assumes a uniform prior on *P* (*model*) and performs a statistical model selection based on the principle of minimum description length (Methods). In general terms, this class of algorithms seeks the most compact possible representation of the data.

In the context of single-cell omics data, the observations correspond to molecular measurements, such as scRNA-seq or scATAC-seq matrices [Fig. 1 A], and the model is a hierarchy of partitions of both molecular features and cells. In previous works, hSBM was extended, as mentioned before, to nSBM[16] to accommodate a data structure containing multiple sources. Now, the bionSBM algorithm is efficiently implemented to handle a multipartite network built from single-cell data, where nodes represent different entities, such as genes, peaks, or proteins, and cells [Fig. 1 B]. Genes and peaks are connected to cells based on the levels of their molecular measurements, using weighted edges. bionSBM allows multiple feature types to be associated with the same data point (i.e. cell) [Fig. 1 B]. This enables the concurrent modelling of multiple modalities in a single inference run [Fig. 1 B, C]. bionSBM is integrated into the single-cell Python universe[20] by handling AnnData[21], for single modality data, and Muon[22] objects for multi-modal data.

In addition, bionSBM distinguishes itself from nSBM in terms of time and scalability by efficiently creating large sparse graphs, which are characteristic of single-cell experiments [SuppFig. 1 A, B]. A key advantage of bionSBM over other algorithms is that, since it operates on graphs, preprocessing does not require scaling or harmonizing (i.e. batch correction) values across experiments. Moreover, features with heterogeneous statistical characteristics (counts, continuous values, or binary measurements), originating from different omic layers and preprocessing pipelines, can be incorporated directly into the multipartite graph, as the inference procedure does not require cross-modality scaling or harmonization. The output of bionSBM is a hierarchical clustering of cells, while providing modality-specific topic assignments [Fig. 1 D]. In other words, the terms “clusters” and “topics” assume a different meaning than in LDA-based approaches: here, a cluster is a set of cells, like in a community detection method, but a topic is a set of features. This allows the creation of groups of cells (clusters) [Fig. 1 C], groups of peaks (peaks-topics) [Fig. 1 C], groups of genes (gene-topics) [Fig. 1 C], or any other kind of feature, for the same multipartite graph. This richer representation allows the disentangling of distinct feature sets across modalities. It is important to note that bionSBM requires that nodes of a given type be non-mixed, so that blocks contain only homogeneous nodes; hence, each group will contain either cells, genes, peaks, etc.

### Model benchmarking

We compared the performance and results of bionSBM with those of ShareTopic[18] and Mowgli[19], the two main topic modelling algorithms applied to multi-modal single-cell data. We compared the performance of these methods on six different data sets. We chose them to represent a variety of situations, including mouse and human genomes, from fully developed to differentiating cells. The data sets show a heterogeneous complexity, ranging from a few thousand to forty thousand cells and from ten to thirty-five cell types. The number of genes per experiment is one of the most homogeneous characteristics across datasets, because the species fix it [SuppFig. 2]. Four of the datasets considered (PBMC from 10x database, BMMCMultiOme[23], HSPC[24], MouseSkin[3]) consist of gene expression and chromatin openness (scRNA-seq and scATAC-seq). From these, the MouseSkin data set has been profiled using the SHARE-seq technology[3], whereas the other three are based on the 10X Multiome platform. Two more data sets (Spleen[25] and BMMCCite[26]) are CITE-seq experiments that combine gene expression and surface protein data. This ensemble of different data was chosen to test the efficacy of the method across different contexts, ensuring that performance is not driven by technological, species- or -omic biases (see SuppTable1, Methods).

For the benchmarking, ShareTopic can handle only scATAC-seq and scRNA-seq pairs; therefore, for the two Cite-seq data sets, we will compare bionSBM only with Mowgli. For the other four datasets, we could compare all three algorithms. For each data set, we explored different options for inputting transcriptional features. In a previous study[17], we showed that cell classification improves when analysing the expression of protein-coding genes and lncRNAs as separate omics layers using the aforementioned nSBM approach. Hence, we created three transcriptomic count matrices: the total Gene Expression (GEX), which contains all the possible transcripts measured by a scRNA-seq experiment; the “mRNA” matrix, containing only protein-coding genes; and the one named “lncRNAs”, which consists only of long-non-coding RNAs.

We tested the quality of the results on the cell layer separately (evaluating the algorithm’s ability to identify cell types) and on the feature layers (evaluating the quality of the topics across the different feature spaces). For each test setting, we ran the algorithms multiple times to explore the stochasticity of the results and the impact of different combinations of transcriptional features. bionSBM, ShareTopic, and Mowgli were executed 25 times per data set and feature-space combination, and we measured running time and memory usage to provide an accurate report of the required resources [SuppFig. 1 C, D].

All the methods embed the input spaces into a low-dimensional representation. Mowgli and ShareTopic create a single low-dimensional topic space, where each topic is a mixture of features from all omics, making interpretation more laborious because different omic features are combined by the algorithm. Instead, bioSBM treats each -omic separately in its own output. Therefore, when using the term “topic”, for bionSBM we will specify what -omic we refer to (mRNA-topic, peaks-topic, etc), while for ShareTopic and Mowgli the topics will be a mixture of them.

### bionSBM better recapitulates cell type in all the designed experiments and across technologies

We first evaluated the performance of bionSBM, ShareTopic, and Mowgli in identifying different cell types across the data sets. Both Mowgli and ShareTopic require the number of desired topics as input (one of their hyperparameters), whereas bionSBM automatically infers it. The same applies to the number of clusters. In this work, for comparability, the number of topics in ShareTopic and Mowgli was set to match the number of annotated cell types, constraining the model to the expected biological heterogeneity. Notably, varying the number of clusters had little impact on performance [SuppFig. 3].

To measure the agreement between the clusters obtained with the three algorithms and the ground truth annotations for the six data sets, we computed the logarithm of the Normalised Mutual Information NMI/NMI* (Methods). The larger the NMI/NMI* ratio, the better the agreement. Since this measure compares the clustering results to a random assignment of cells in clusters, it is affected by the characteristics of the specific data set, and the results should not be used to compare data sets (for instance, the number of cell types is very different across data sets). bionSBM consistently outperforms Mowgli for the two CITE-seq data sets across all combinations of omics [Fig. 2 A, D]. For the scATAC-seq + scRNA-seq data sets, the three algorithms show, in general, similar performances in the BMMCMultiOme and HSPC data sets [Fig. 2 E, F], while for the PBMC and MouseSkin data [Fig. 2 B, C], bionSBM performs better than the other two algorithms. It is interesting to notice that the performance of bionSBM is particularly good on the three data sets with the highest number of cell types [Fig. 2 A, C, D], reflecting superior performance on more complex data.

**Figure 2.**
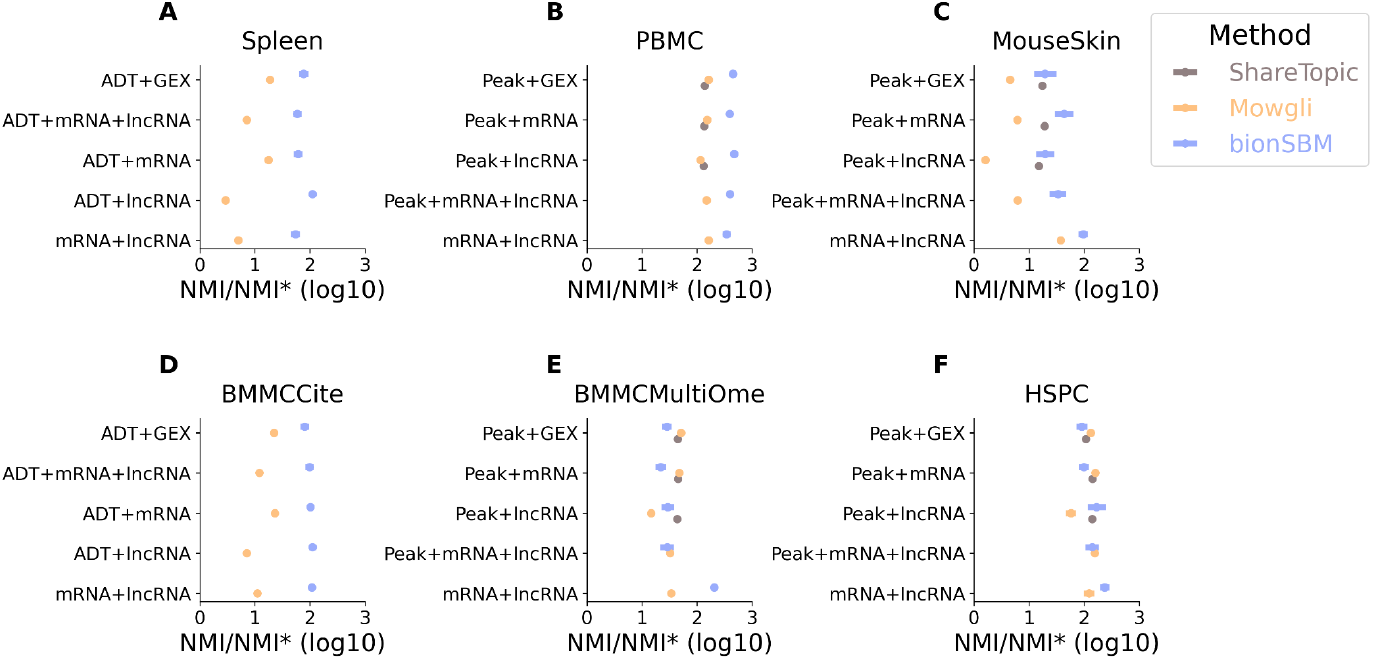
NMI/NMI* score between each algorithm partition and ground truth annotation for the six data sets. Each dot represents the mean value, and the error bar represents three times the uncertainty on the mean. The higher the values, the closer the clustering is to the ground truth annotations.

### Graph based topic modelling signatures are highly specific and clearly distinct

An important feature of algorithms belonging to the framework of topic modelling is their probabilistic output. Using the *P* (*topic* | *cell*) probability distribution one can associate to each cell the most relevant topic, i.e. the one with the highest *P* (*topic* | *cell*), and then aggregate them per cell type or cluster. Moreover, one can assess the quality of this topic assignment using two standard metrics: specificity and distinctiveness. The former metric measures how group-specific the dominant topic is (i.e. the topic with the highest probability), while the latter captures how well topics are separated by quantifying the in-group topic probability and the out-of-group topic probability (Methods). Both metrics take values in the range [0-1], with 1 being the maximum performance. We will assess them for all three topic modeling methods using the predefined cell types as groups. As mentioned above, bionSBM embeds in separate spaces each different feature (mRNAs, lncRNAs, ATAC-seq peaks, etc.), whereas Mowgli and ShareTopic create a set of topics which are a mixture of features. Hence, the specificity and distinctiveness of bionSBM is computed for each -omic space, while for the other methods the topic space is mixed.

bionSBM achieves, in general, a high topic specificity [Fig. 3], higher than the other methods. The specificity is high in all datasets and modalities, meaning that the method is robustly able to explain every cell type from each modality. In agreement with the classification results [Fig. 2], bionSBM achieves a particularly high specificity in the Spleen and BMMCite data sets [Fig. 3 A, D], which are the most complex ones in terms of cell types [SuppTable1]. The three algorithms achieve similar values for all the data sets in their distinctiveness metric [Fig. 4], with good performance in general, around 0.75.

**Figure 3.**
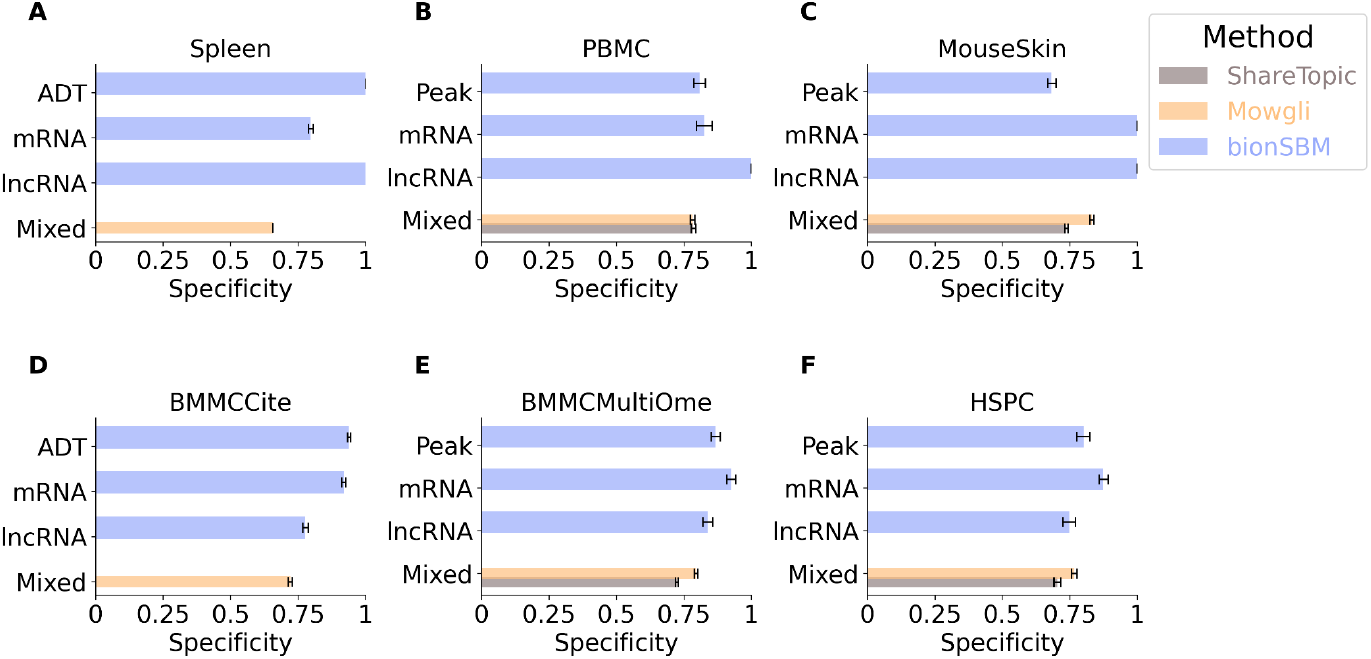
specificity (the measure of how cluster-specific the dominant topic is) of the topics for the six data sets. Each bar represents the mean value over multiple runs and the error bar represents three times the uncertainty on the mean.

**Figure 4.**
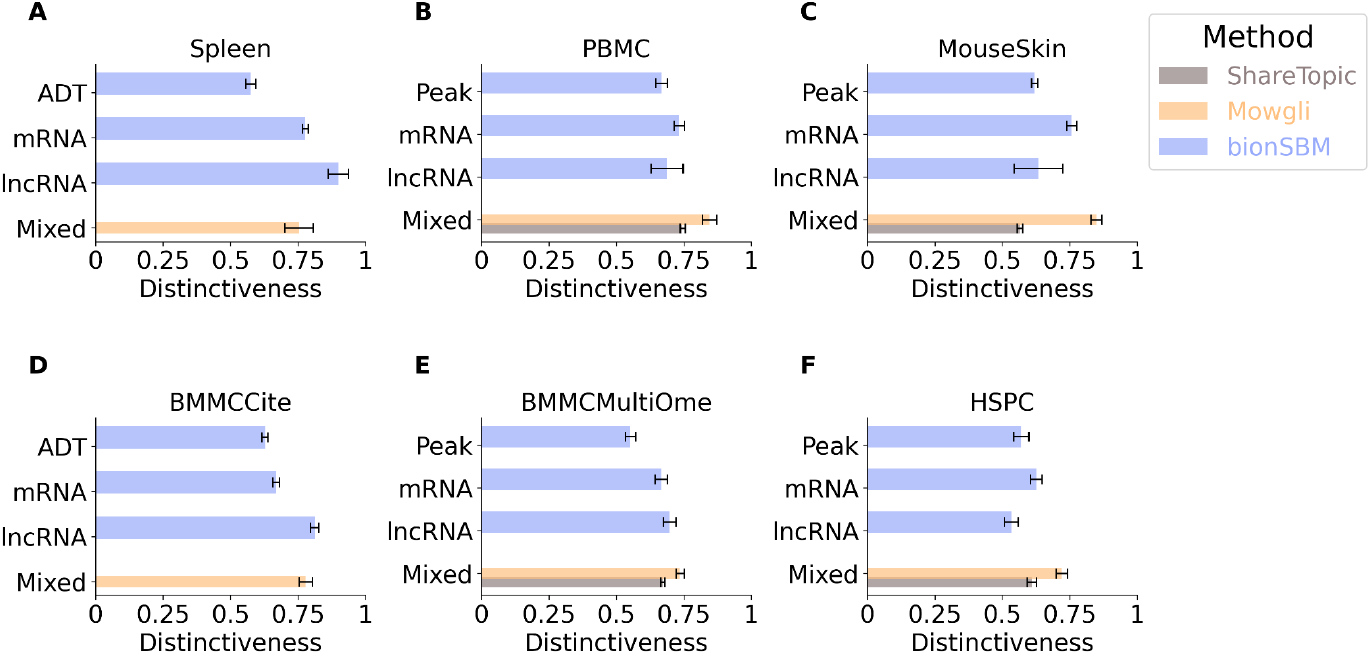
distinctiveness (measure of how well topics are separated by quantifying the in-cluster topic probability and the out-of-cluster probability) of the topics for the six data sets. Each bar represents the mean value over multiple runs and the error bar represents three times the uncertainty on the mean.

Together, these two metric results show that bionSBM’s signatures are highly specific and show strong in-group enrichment relative to out-of-group probability; thus they can be used to assign a biological explanation to the cell type groups, making the method easy to interpret and translational. All three methods achieve similar distinctiveness results, meaning that all topic models analysed assign distinct topics per cell type, but bionSBM outperforms the others in the specificity, meaning that it tends to assign topics to a cluster in a more specific way.

To biologically interpret the inferred topics, we assigned a topic to each cell type using the following criterion: for mRNA and peak modalities, we computed *P*_*c*_ (*topic* | *cell type*) and assigned to each cell type the topic with the highest distinctiveness (Methods). We then examined the top-ranked genes by *P* (*gane* | *topic*) in the selected mRNA topic and compared them with transcription factor binding motifs enriched in the corresponding peak topic for the same cell type obtained with HOMER[27]. Linking the chromatin-topics to the transcriptome-topics we could extract meaningful case-studies that substantiate the interpretability of bionSBM.

This approach recovered well-established regulatory programs across data sets. In PBMCs, the peak-topic with the highest distinctiveness for the B memory cell population was enriched for the PAX5 motif [Fig. 5 A], an essential gene required for B cell commitment from lymphoid progenitors[28]. In a coherent manner, the same topic was also the highest enriched in B naive cells [SuppFig. 4 A], and the same gene was found to be in the most enriched mRNA-topic in both cell states [Fig. 5A, SuppFig. 4A]. Notably, the topics were highly enriched for both the mRNA and ATAC features only for the B cell type populations [SuppFig. 5 A], highlighting the specificity of our approach. In the HSPC data set, granulocyte monocyte progenitors (GMPs) had the highest association with a peak-topic showing the motif of IRF8 [Fig. 5 B], a TF known to regulate the differentiation of GMP cells[29]. The corresponding highest associated mRNA-topic for the same cell type contained the same TF [Fig. 5 B], again connecting the two feature spaces mechanistically. Lastly, in the BMMCMultiOme data set, the proerythroblasts, normoblasts, and erythroblasts had a high probability for a peak-topic showing the motif of KLF1 [Fig. 5 C, SuppFig. 4 B, SuppFig. 4 C], a master regulator of erythropoiesis[30], and the same gene belonged to the most relevant mRNA-topic for the three above mentioned cell types, which are the primary cell states evolving during erythropoiesis [Fig. 5 C, SuppFig. 4 B, SuppFig. 4 C]. Again, the selected mRNA- and peak-topics were not only the ones with the highest distinctiveness among all other topics, but they were also highly enriched only for the cell types under examination [SuppFig. 5], giving evidence that the explanations are specific.

**Figure 5.**
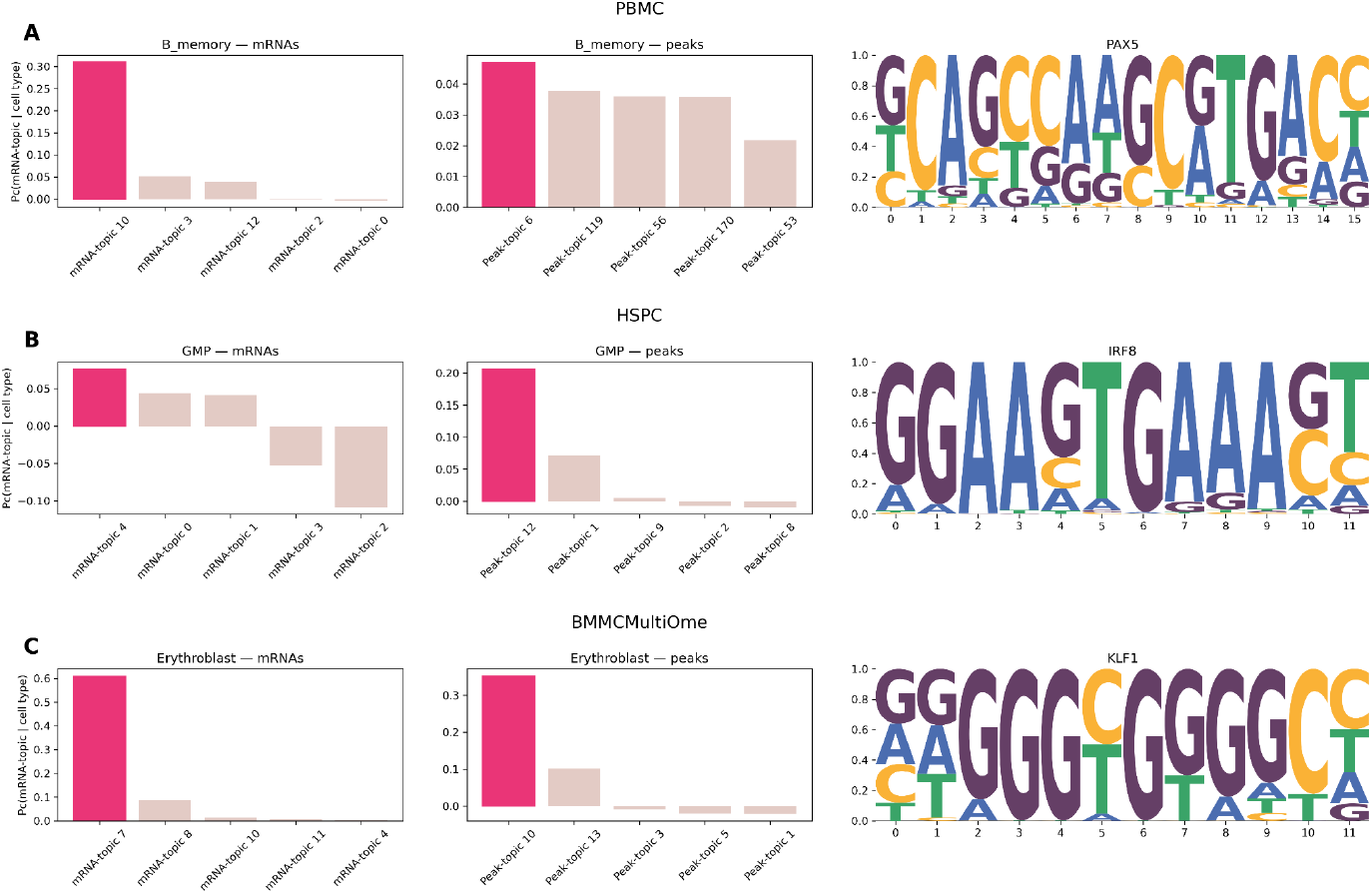
for each data set, three panels are shown. **Left:** enrichment profile of the Peak-topic with highest distinctiveness for the given cell type. **Middle:** enrichment profile of the mRNA-topic with highest distinctiveness for the given cell type. **Right:** sequence logo of a significantly enriched transcription factor motif identified in peaks belonging to the selected peak-topic (HOMER analysis).

Overall, the discovered peak and mRNA assignments consistently recovered canonical lineage-defining transcription factors, supporting the ability of the bionSBM integrative topic model to capture biologically meaningful, cell–type–specific regulatory programs across modalities and to link the transcriptome to the epigenome.

## Discussion

In this study, we present bionSBM, a graph-based stochastic block model to integrate single-cell multi-omic data [Fig. 1]. This approach has both theoretical and practical advantages over the state-of-the-art topic models for the single-cell field. First, our method can be applied to any multi-modal data since it can handle an arbitrary number of partitions (i.e. modalities). Second, the method automatically detects the optimal number of topics and clusters without the need to specify these values as hyperparameters. Moreover, all molecular layers are equally weighted, preventing the domination of one -omic due to technical differences that are not solvable with normalisation approaches. Finally, because topic modelling algorithms are Bayesian, they require specification of a prior distribution over topics. Rather than imposing a structured distribution that constrains cells and features to follow a single mixture model, we employ a flat (non-informative) prior.

Integrating multiple omics layers is key to uncovering new biological insights. By modelling separate topic compositions for each network layer, bionSBM captures coherent relationships across modalities, inferring modality-specific topic structures within a unified probabilistic framework, enabling independent yet coherent biological interpretations across omic layers.

bionSBM also improves performance in cell type identification, outperforming state-of-the-art approaches and more accurately recovering known biological annotations [Fig. 2]. These performance gains come along with inherent interpretability. The model generates specific [Fig. 3], distinct [Fig. 4], and stable [SuppFig. 6] topic representations that provide biologically meaningful explanations across molecular layers [Fig. 5, SuppFig. 5, SuppFig. 6]. This balance between accuracy and interpretability is essential, as predictive improvements alone are unlikely to drive meaningful biological discovery.

We released a Python implementation of the tool, integrated into the single-cell universe, leveraging the convenient AnnData and Muon packages, guaranteeing reproducibility and broad usability of the method. Here, we have applied bionSBM to single-cell multiomic measurements from scRNA-seq, scATAC-seq, and scRNA-seq with single-cell protein measurements (CITE-seq). However, the framework can be applied to any new omic measurements that can be summarised as a count matrix, making the approach truly generalizable.

## Methods

### Single-cell data processing

The raw reads of the PBMC data set were processed using Cell Ranger Arc 2.0.2, aligning the reads onto the complete human genome (T2T)[31]. The gene expression count matrices of all the other data sets (BMMCMultiOme[23], BMMCCite[26], HSPC[24], mouse skin[3], spleen[25]) were taken from the literature together with the cell type annotations. From the total GEX count matrices, which contain all the families of transcripts that scRNA-seq can profile, we extracted the mRNAs and lncRNAs counts using the genome annotations from Esembl[32]. All the matrices followed the same preprocessing according to the standard single-cell best practices[5]: quality control, filtering, library size normalisation and log transformation.

The PBMC peaks count matrix has been built with Episcanpy[33] after peak calling with MACS2[34]. All the other peak count matrices have been obtained from the literature. Similar to scRNA-seq data, the preprocessing followed the best practices for scATAC-seq data with quality control, filtering and library size normalisation. In both scRNA-seq and scRNA-seq matrices, we kept only the highly variable features. The ADT (surface protein) count matrices have been only library size normalised because the number of features is extremely limited, so no additional filters are needed.

### Stochastic block models

The statistical inference procedure, as well as the definition of topics and probability distributions, is based on the hSBM algorithm. The input matrices are transformed into a weighted multipartite network. Each node represents a feature or a cell (i.e. mRNA, lncRNA, peak ADT) and the feature nodes are connected to the cell node with an edge whose length is proportional to the expression of the gene in the cell or the openness of the peak in the cell. The algorithm defines a P(model|data) where the data is the graph and the model is the particular partition built during the inference. The inference process begins with nodes (i.e. genes or cells) assigned to random blocks (either clusters or topics). The process consists of a Markov Chain Monte Carlo (MCMC) simulation to adjust node assignments to blocks to minimize the description length (DL)[35], defined in natural units as the number of bits required to encode the model and the data. Moves in the MCMC are accepted with probability[36]:

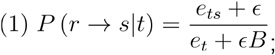

which represents the probability of moving a node from block (cluster or topic) *r* to block *s* if a random neighbor of the considered node belongs to block *t*; given that *B* is the total number of blocks, *e*_*ts*_ is the number of edges from group *t* to group *s* and *e*_*t*_ is the total number of edges within group *t*. Such probability is proportional to the ratio between the number of edges that connect group *t* and block *s* and the number of edges that go to block *t* to group *s* and is *e*_*t*_ the total number of edges within group *t*. Such probability is proportional to the ratio between the number of edges that connect group *t* and block *s* and the number of edges that go to block *s*, plus a term that is necessary in order to have a uniform probability if there are no connections between the blocks considered. For instance, if all the edges from *t* go to group *s, e*_*ts*_ equals *e*_*t*_ and the probability becomes uniform: 1/*B*. The algorithm supports multiple random initializations, i.e., different starting points for estimating the posterior probability.

It is important to highlight that, looking at the probability of accepting a move, the advantage of using multibranch SBM on multi-omics problems is clear: when moving a cell from block *r* to block *s*, the algorithm samples a neighbor of the cell and it labels its group as *t*. At this point, the model only considers the links between the cell and the nodes in the group, so the normalisations of other branches are irrelevant. The final output of the model consists in a hierarchy of topics and clusters. For each level in the hierarchy, the model returns the grouping of features into topics, of cells into clusters and the probability of the topic to belong to each cell[12,13,16]. A first implementation of multipartite hSBM, the so-called nSBM, is available at https://github.com/BioPhys-Turin/nsbm[16][37,38] based on the graph-tool library[39].

### Experiment running and metrics

bionSBM has been run 25 times with each combination of feature spaces changing the initial seed every time and setting the number of initialisation to 7 as suggested in [17]. ShareTopic and Mowgly experiments have been run 25 times as well. In Mowgli, we set the strength of the regularisation to 0.05 for all the feature spaces to treat them equally. For each outcome of the experiments, and for each level of bionSBM hierarchy, we have measured the NMI/NMI* between the clustering of the algorithms and the original annotation of the cells in cell type.

The NMI/NMI* is a version of the normalised mutual information (NMI)[40] that accounts for randomness, where NMI* quantifies the agreement between the ground truth label and a random clusterization of the cells. In this way, we account for partitions with a massive number of clusters that would falsely lead to high NMI. On a log scale, the lower limit is 0, indicating the clustering has been totally dominated by randomness, i.e. the NMI is equal to the NMI*. We kept the bionSBM level with the highest NMI/NMI* and averaged the results for each method and combination of feature spaces to produce the plots of Fig. 2.

### Topics specificity, distinctiveness and stability

To compute the specificity, distinctiveness and stability we selected, for each experiment and for each run, the level with the closest number of clusters to the ground truth annotation. After setting the level, we assigned to each group of cells the topic with the highest probability. We used a modified version of the *P* (*topic* | *cell*) defined as *P*_*c*_ (*topic* | *cell*) (where c stands for centered) to exclude topics that are important for all the cells which would be the equivalent of the topic of articles and prepositions in NLP, as suggested in[12]. The centred probability is defined as

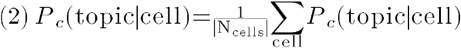

From which we can derive the same quantity for a specific group of cells as

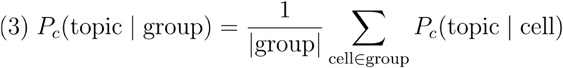

We then computed how many different topics have been assigned to the groups of cells and divided that number by the total number of cells groups. We select the level of the hierarchy in bionSBM that has the closest number of topics to Mowgli and ShareTopic. We average the values obtained from the runs of the tools to produce the bar plot of Fig. 3.

Once we marked the most important topic to each group as described above, we quantify the distinctiveness of these assigned topics, which is defined as

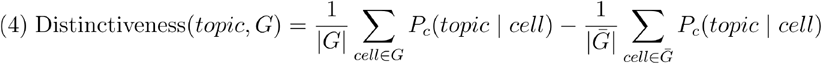

where *t* is the topic assigned to group *G, P*_*c*_ (*t* | *c*) is defined as in (1) and 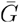 is the set of all the cells expect the ones in group *G*. In other words, it is defined as the difference between the average probability of the topic belonging to the cells within the group and the average probability of belonging to the cells out of the group. Like for the specificity, we average the values across the multiple runs of the model to produce Fig. 4.

To quantify the stability of topic–feature associations across independent runs of the same experiment, we measured the similarity of the top three most important topics identified for each cell type. For a given run *r*, let *P*_*c,r*_ (*t* | *c*)denote the centred posterior probability of topic *t* for cell *c*, as obtained from the topic–document matrix as shown in Eq 2.

For each cell type τ, a cell-type–specific topic score was computed by averaging centred probabilities across cells of that type:

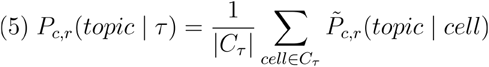

where *C* _τ_ ⊂ *C*is the subset of cells annotated as cell type τ. For each run and cell type, the set 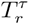 of the top *k* = 3topics with the highest values of *P*_*c,r*_ (*topic* | τ)was selected. Each topic 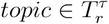 is associated with a topic–feature probability vector *Z*_*r*_.( · | *topic*)

To compare two runs *r*_*i*_ and *r*_*j*_, we computed pairwise similarities between all topic vectors in 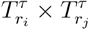 using the cosine similarity,

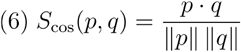

where *p* and *q* are normalized topics–features distributions.

For each cell type and run pair, the similarity was defined as the maximum similarity across possible combinations given the three top topics for run pairs:

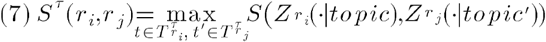

The choice of taking the three top topics makes the metric robust to slightly different ranking in the top positions of the topics which are due to the noise of the data and the approximation of the sampling procedure. The final stability score for an experiment was obtained by averaging *S* ^τ^ (*r*_*i*_, *r*_*j*_) across all cell types and all unordered pairs of runs.

## Supplementary figures

**Supplementary figure 1.**
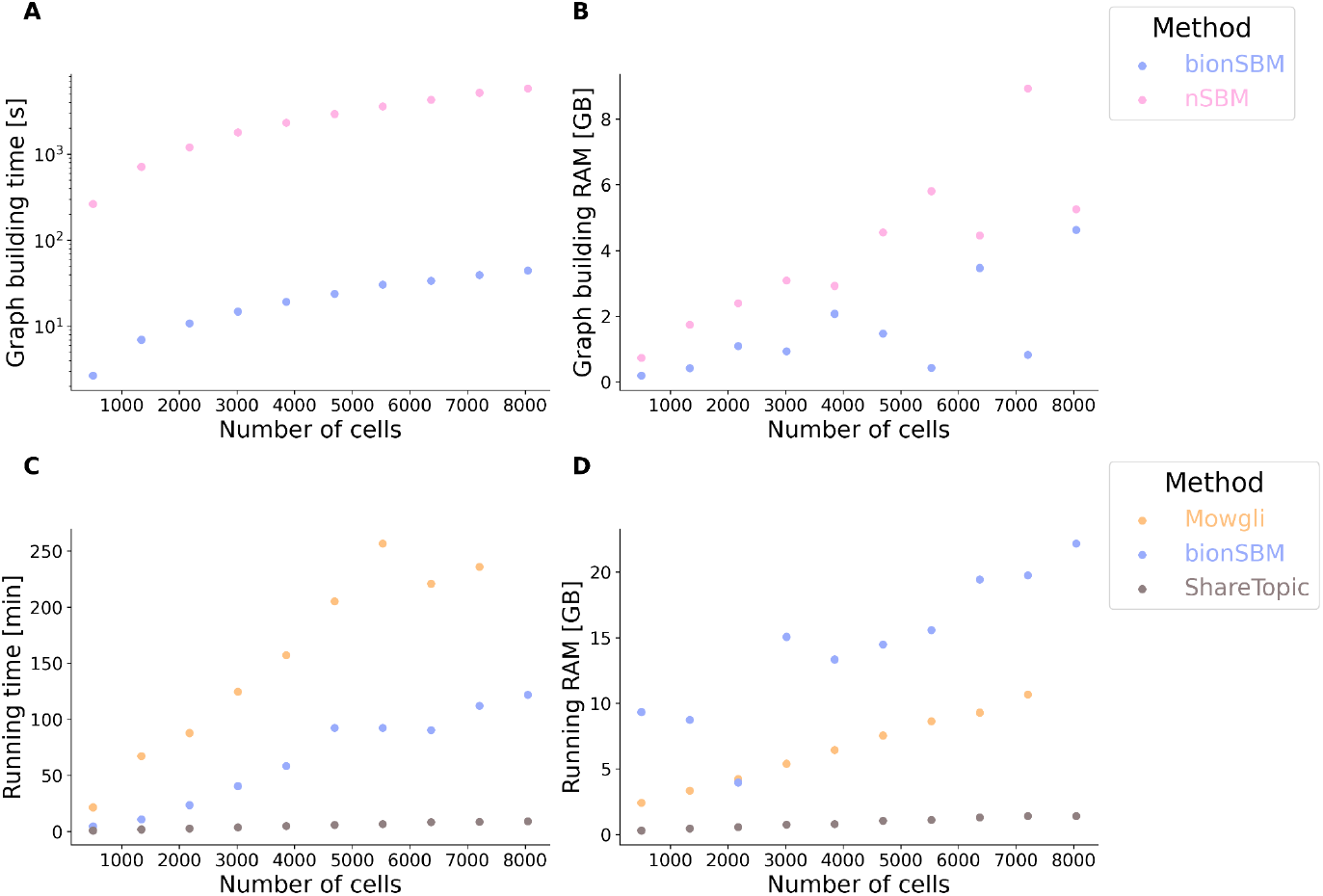
**A-B** performances varying the number of cells of bionSBM and in creating the graph derived from the count matrix. The downsampling is stratified by cell type abundance to preserve the biological structure of the data set. The running time (**C**) and memory requirements (**D**) of the three topic modelling methods vary with the number of cells.

**Supplementary figure 2.**
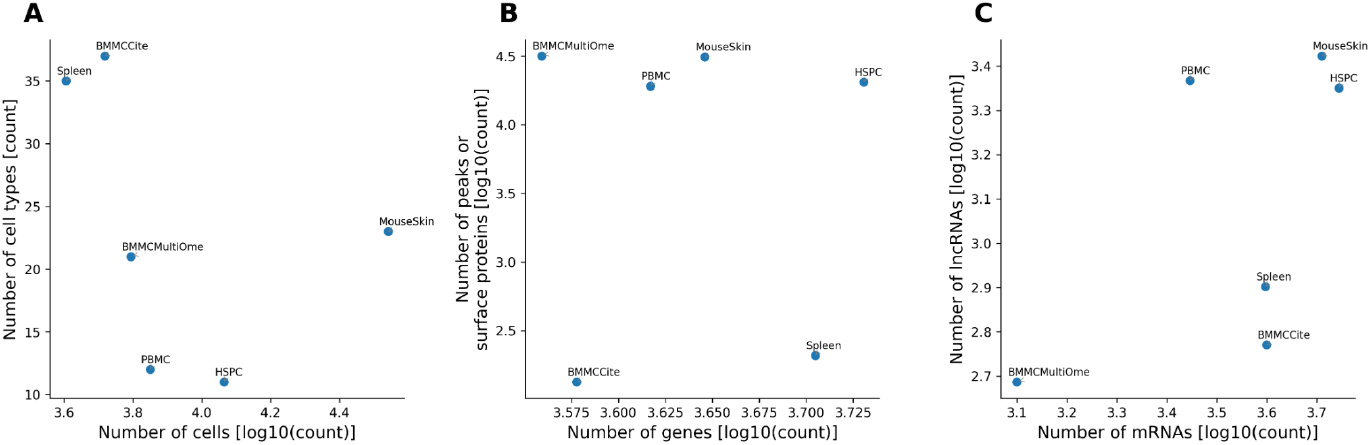
complexity of the six data sets measured as the number of cells and the number of features (genes, peaks).

**Supplementary figure 3.**
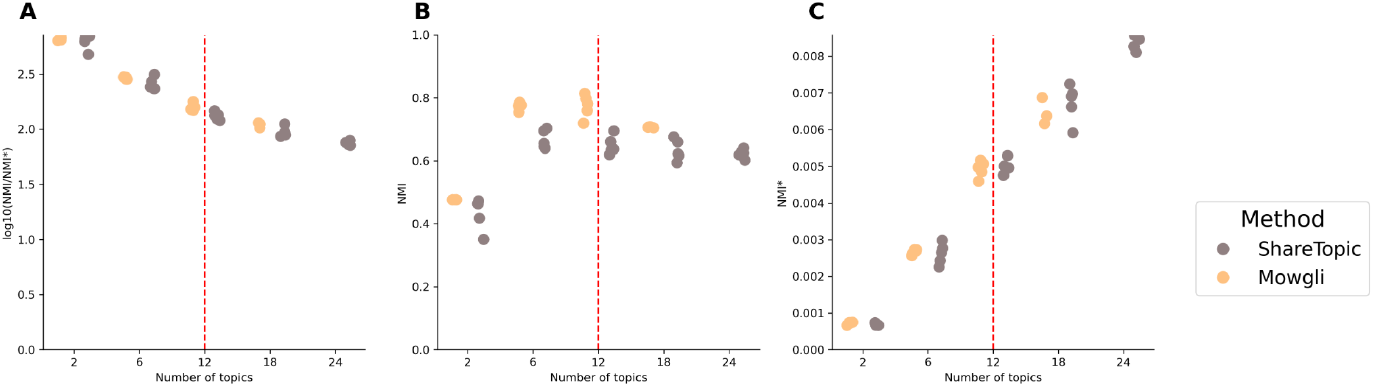
trend of NMI/NMI* (**A**), NMI (**B**) and NMI* (**C**) of Mowgli and ShareTopic varying the number of required topics for the PBMC data set. The vertical dashed line highlights the real number of cell types. The combined metric (A) slowly decreases with the the number of topics due to the increase in NMI*, as expected (the residual entropy is higher in a finer partition). The values of the NMIs are stable (B). These results suggest that performance is not strongly dependent on the number of topics.

**Supplementary figure 4.**
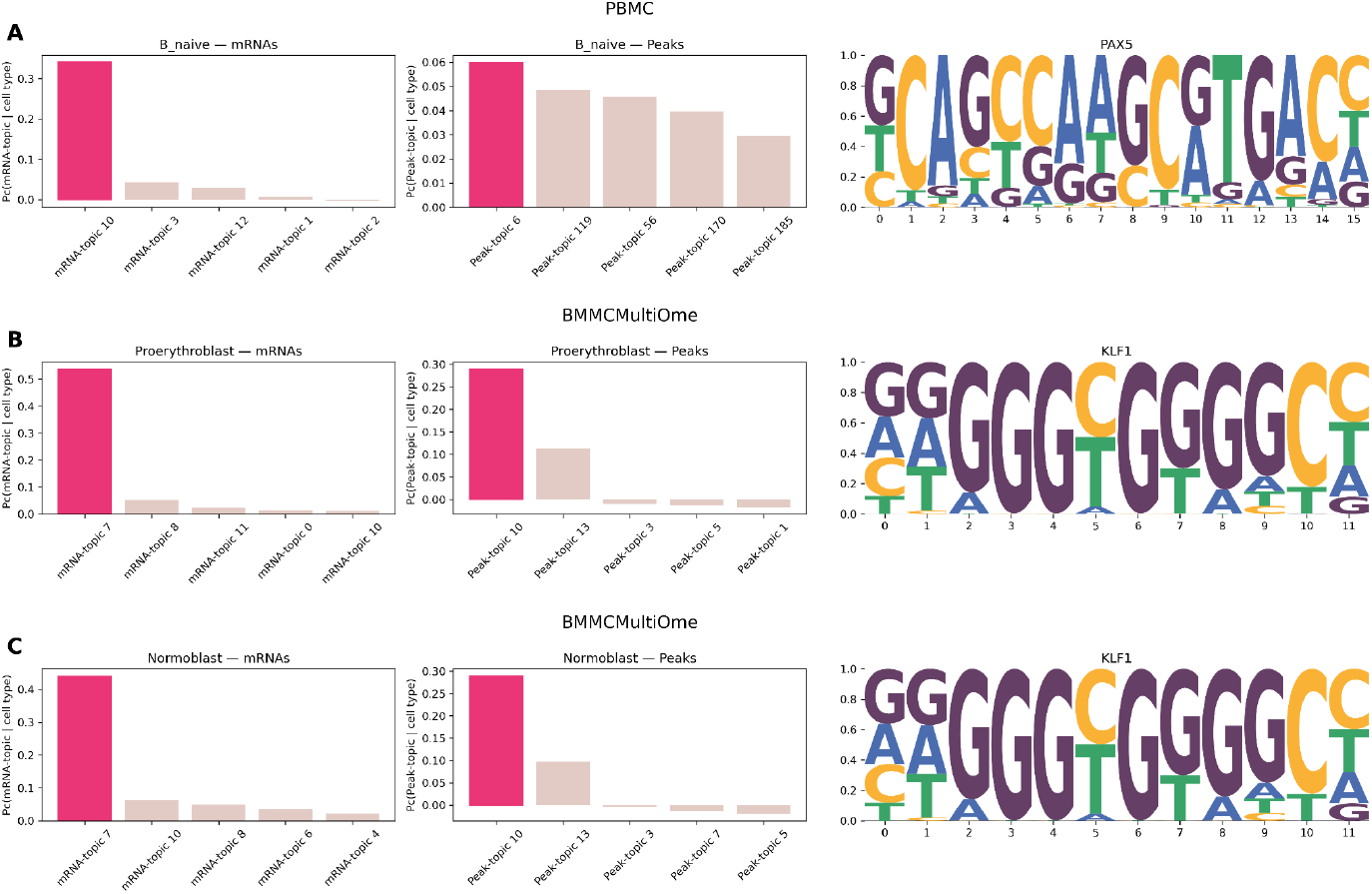
for each data set, three panels are shown. **Left:** enrichment profile of the Peak-topic with highest distinctiveness for the given cell type. **Middle:** enrichment profile of the mRNA-topic with highest distinctiveness for the given cell type. **Right:** sequence logo of a significantly enriched transcription factor motif identified in peaks belonging to the selected peak-topic (HOMER analysis).

**Supplementary figure 5.**
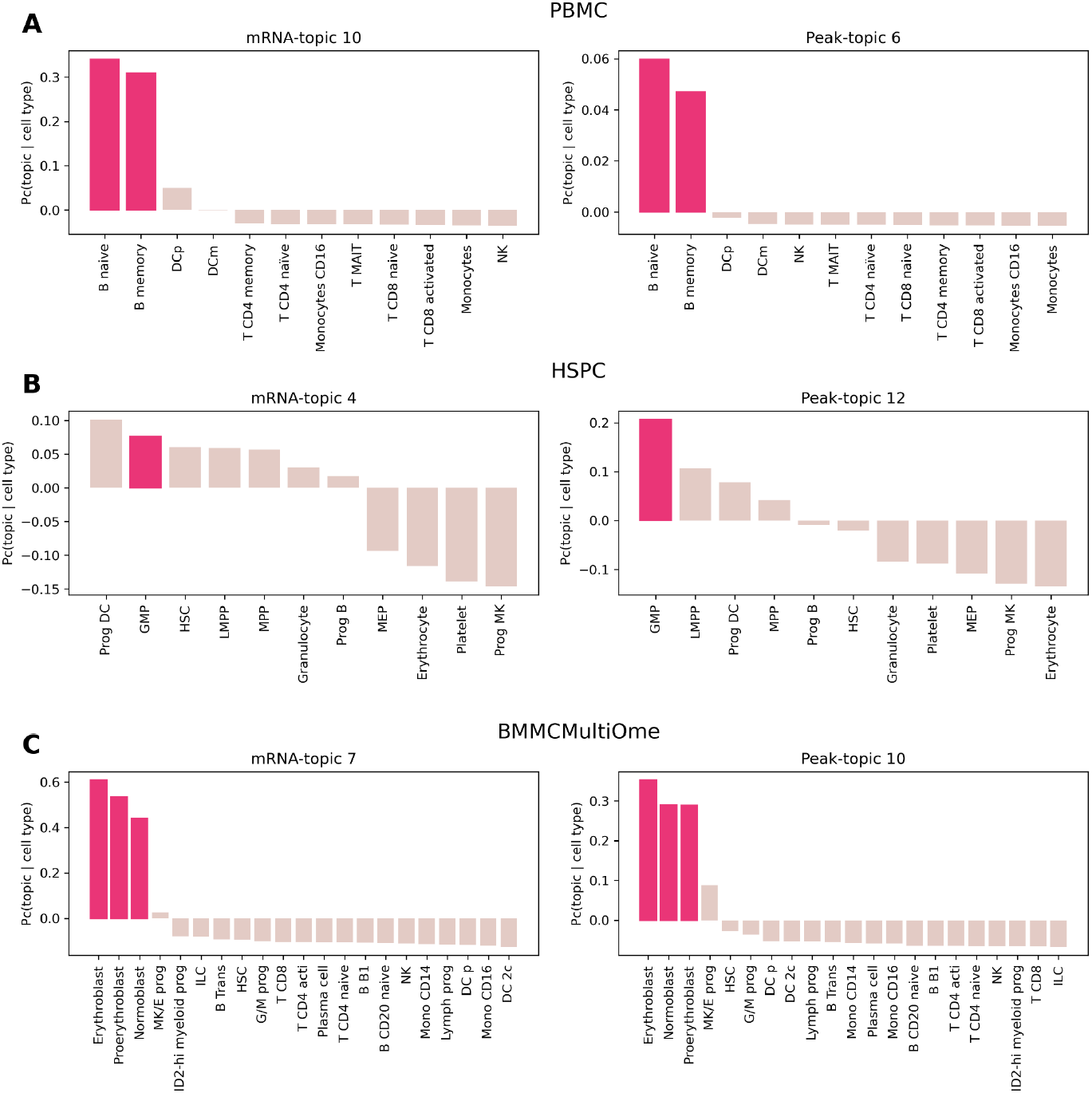
for each dataset, two panels are displayed. **Left:** enrichment profile of the mRNA-topic with the highest distinctiveness for the selected cell type, shown across all other cell types. **Right:** enrichment profile of the Peak-topic with the highest distinctiveness for the selected cell type, shown across all other cell types.

**Supplementary figure 6.**
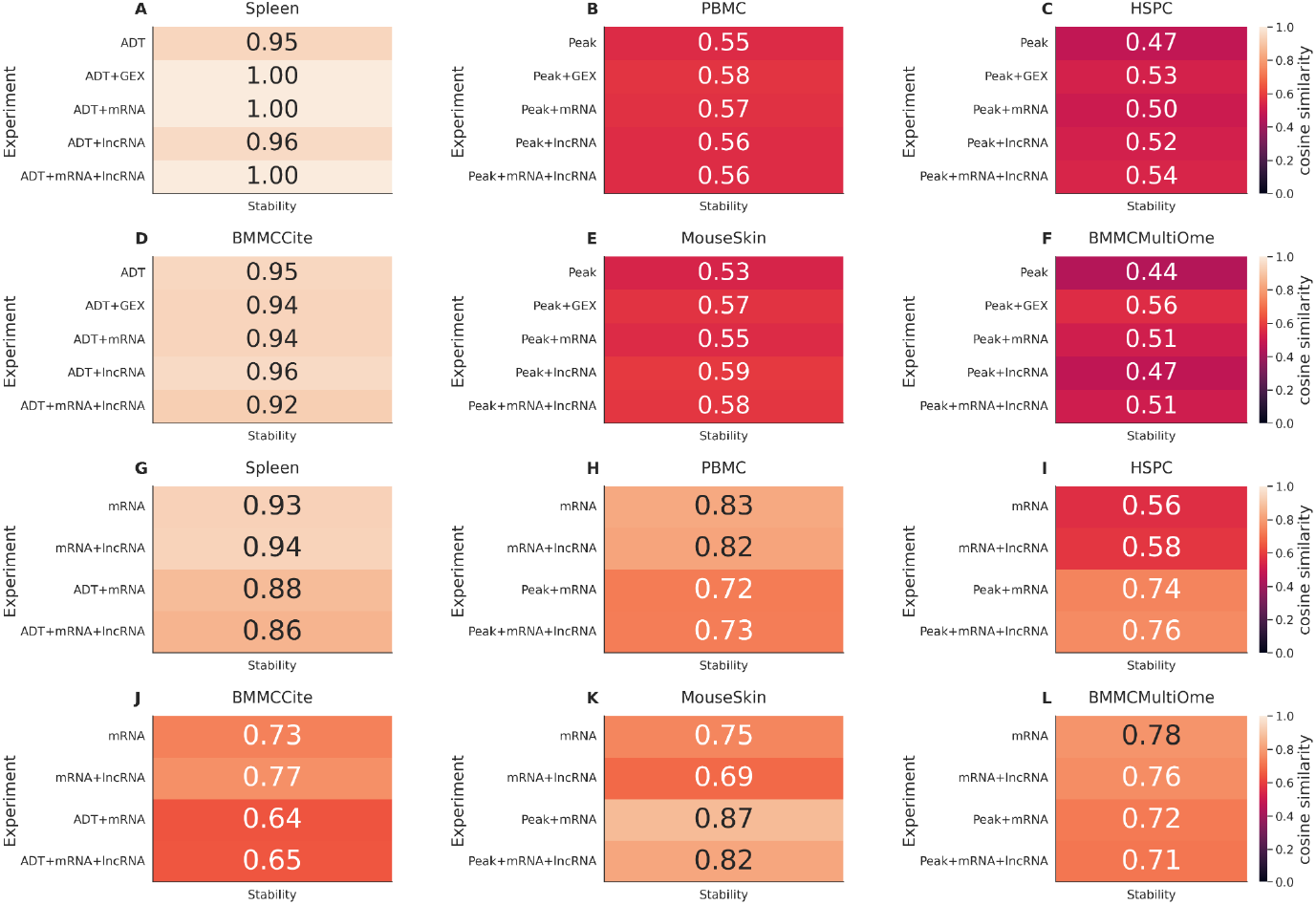
stability of the bionSBM algorithm. Average cosine similarity across 25 runs of the same experiment of the three most important ADT (**A,D**), peak (**B,C,E,F**) and mRNA (**G-L**) topics for each cell type.

## Supplementary tables

**Suppl Table 1.**
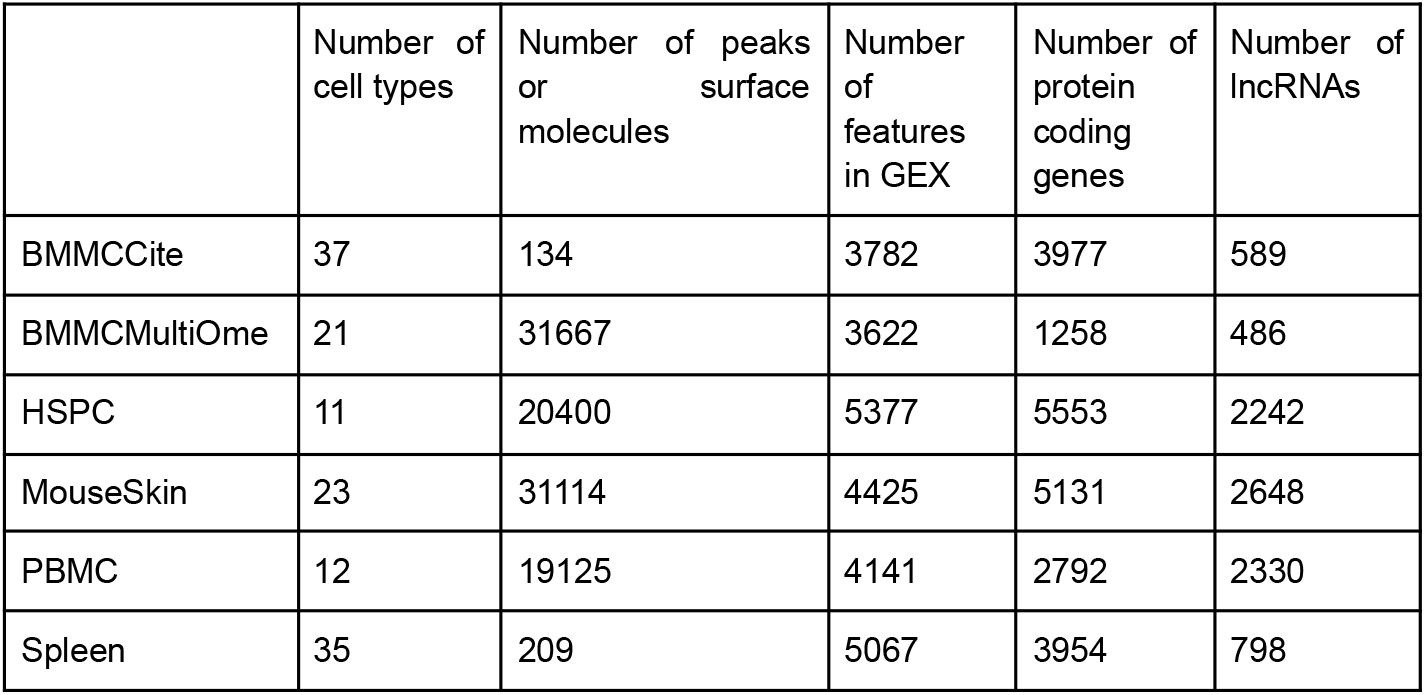
Metadata and properties of the data sets considered in this study.

## Availability of Data and Materials

Code and tutorial for bionSBM are available at https://github.com/gmalagol10/bionsbm.git. The code to reproduce all the results are available at https://github.com/gmalagol10/bionsbm/tree/main/reproducibility.

## Contributions

G.M., F.P., M.C., and M.C.T. designed the study and conceived the algorithm. G.M. implemented the algorithm and performed the benchmarking analyses. F.P. contributed code and assisted with implementation. All authors contributed to the interpretation of the results. G.M., F.P., L.M., M.C., and M.C.T. drafted the manuscript, with contributions from A.T. and A.M. All authors critically revised the manuscript and approved the final version.

## Acknowledgments

G.M. is supported by the Helmholtz International Lab Causal Cell Dynamics (InterLabs-0029) - Grant support from the Initiative and Networking Fund of the Hermann von Helmholtz-Association Deutscher Forschungszentren e.V.-. G.M is also supported by the Helmholtz Association under the joint research school “Munich School for Data Science — MUDS and by German Research Foundation project ID 213249687–SFB 1064.

We thank the BMC Bioinformatics Core Facility for providing access to their HPC cluster. Figure 1 has been made with the help of Biorender[41].

## Ethics declarations

## Ethics approval and consent to participate

Not applicable.

## Consent for publication

Not applicable.

